# Genome-resolved transcriptomics reveals novel organohalide-respiring bacteria from Aarhus Bay sediments

**DOI:** 10.1101/2023.04.17.537210

**Authors:** Chen Zhang, Tom N.P. Bosma, Siavash Atashgahi, Hauke Smidt

## Abstract

Organohalide-respiring bacteria (OHRB) are keystone microbes in bioremediation of sites contaminated with organohalides and in natural halogen cycling. Known OHRB belong to distinct genera within the phyla *Chloroflexota*, *Proteobacteria* and *Firmicutes*, whereas information about novel OHRB mediating natural halogen cycling remains scarce. In this study, we applied a genome-resolved transcriptomic approach to characterize the identity and activity of OHRB from PCE-respiring cultures previously enriched from sediments of Aarhus Bay. Combining short- and long-read sequencing approaches, we assembled 37 high quality bins with over 75 % completeness and less than 5 % contamination. Sixteen bins harbored RDase genes, and were affiliated taxonomically to the class of *Bacilli*, and phyla of *Bacteroidota*, *Synergistota*, and *Spirochaetota*, that have not been reported to catalyze reductive dehalogenation. Among the 16 bins, bin.26, phylogenetically closely related to the genus *Vulcanibacillus*, contained an unprecedented 97 RDase genes. Of these, 84 RDase genes of bin.26 were transcribed during PCE dechlorination in addition to RDase genes from members of *Synergistales* (bin.15 and bin.32) and *Bacteroidales* (bin.18 and bin.24). Moreover, metatranscriptome analysis suggested the RDase genes were likely under the regulation of transcriptional regulators not previously associated with OHR, such as HrcA and SigW, which are known to respond to abiotic environmental stresses, such as temperature changes. Combined application of genomic methods enabled us to pinpoint novel OHRB from pristine environments not previously known to mediate reductive dechlorination and to provide evidence towards the diversity, activity and regulation of reductive dehalogenases.

## Introduction

Organohalide respiring bacteria (OHRB) can derive energy for growth from the use of reductive dehalogenation of halogenated compounds as terminal electron accepting process. They employ reductive dehalogenases (RDases) that catalyze the removal of halide(s) from the carbon backbone via respiratory electron transfer (1, 2). In the process of organohalide respiration (OHR), organohalides serve as the terminal electron acceptor for the electrons derived from, for example, hydrogen or lactate, through a membrane-associated electron transport chain (ETC). In addition to RDases known from OHRB, reductive dehalogenation can also be catalyzed by another type of reductive dehalogenase, thiolytic, glutathione-dependent tetrachloro-P-hydroquinone RDase (TPh-RDase) as previously reported for biodegradation of chlorinated compounds (3, 4). In many cases, RDase gene-containing gene clusters in OHRB are composed of genes encoding a transcriptional regulator, the RDase itself, and a potential electron transporter. Thereinto, three regulatory systems of RDase gene clusters have been characterized: CRP (cyclic AMP receptor protein) / FNR (fumarate and nitrate reduction regulator) superfamily, Two-Component Systems (TCS) and MarR (multiple antibiotic resistance regulator) (5, 6). The minimum cassette of RDase gene clusters is usually composed of a gene encoding the catalytical subunit, *rdhA*, and a second gene, *rdhB,* coding for a cognate membrane-anchoring protein B (7–9). Organohalides are present as natural products in marine environments and other high salt environments (10–12), which is why it is interesting to study their role in the geochemical carbon and halogen cycles and their potential eco-physiological importance.

OHR exploration in marine sediments has been a long-term topic for several decades, which was mainly based on culture-dependent and molecular detection methods, such as dilution-to-extinction isolation, marker-gene amplicon sequencing and quantification (13–15). Recent research into the microbial composition of marine sediments, making use of metagenomic techniques and bioinformatics, revealed the presence of microorganisms from various phyla containing genes predicted to code for RDases, including, next to the commonly known phyla, such as *Chloroflexota* and *Firmicutes*, new archaeal phyla, *Lokiarchaeota*, *Thorarchaeota*, and *Heimdallarchaeota* belonging to the proposed *Asgardarchaeota* superphylum (16).

Based on single-cell amplified genomes (SAGs) retrieved from sediment of Aarhus Bay, a putative RDase gene was found in the assembled genome DEH-C3, belonging to the *Dehalococcoidia* (17). Metatranscriptomic data from the surroundings of the initial sampling site revealed a high abundance of *tceA*-like gene transcripts implicating the potential for reductive dehalogenation in this marine sediment from Aarhus Bay (18). Moreover, five *Desulfatiglans*-related SAGs were found bearing putative RDase genes from the sulfate-rich subsurface of Aarhus Bay marine sediments, and it was speculated that OHR might be an alternative energy conservation strategy under sulfate-limiting conditions (19). Nevertheless, although these analyses of extensive DNA and RNA sequence data indicated the existence of potential OHR in marine sediments of Aarhus Bay, unambiguous physiological proof remains pending, and the OHRB involved were not isolated nor unequivocally identified.

In a previous study we have found that marine sediments from Aarhus Bay can dehalogenate a range of halogenated compounds, including tetrachloroethene (PCE), 2,6-dibromophenol (2,6-DBP), 1,4-dibromobenzene (1,4-DBB), 3-bromophenol (3-BP), and 2,4,6-triiodophenol (2,4,6-TIP) (20). 16S ribosomal RNA (rRNA) gene amplicon sequencing revealed that members of the phyla *Desulfobacterota*, *Firmicutes* and *Bacteroidota*, such as *Desulfovibrio*, *Desulfuromusa* and *Bacillus*, were enriched in sediment-free PCE dechlorinating cultures. *Desulfovibrio* and *Desulfuromusa* were recently shown to be capable of reductive debromination of 2,6-DBP to phenol (21). However, *Bacillus* spp. or members of the *Bacteroidota* have never been reported before to use organohalides as terminal electron acceptors.

Thus, the aim of this study was to identify the OHRB involved in the dehalogenation of PCE aligning to the previously reported study (20). To this end, we combined metagenomics and - transcriptomics of sediment-free PCE-dechlorinating enrichment cultures with physiological observations in order to identify the corresponding OHRB and their RDase genes for PCE dechlorination with(out) additional sulfate. This revealed the presence of a large number of genomic bins encoding putative reductive dehalogenases, as well as the expression of many of these genes that appeared to be under the control of transcriptional regulators not previously associated with OHR.

## Materials and methods

### Chemicals

PCE, TCE, cDCE, VC, and ethene were purchased from Sigma-Aldrich. Stock solutions of lactate (0.5 M) and sulfate (0.5 M) were prepared separately by filter sterilization (syringe filter, 0.2 µm, mdimembrane, Ambala Cantt, India). All other (in)organic chemicals were obtained in analytical grade or higher.

### Cultivation

Cultures were incubated in marine medium under sulfate-free (NS) or sulfate-amended (S) conditions as described previously (20). These cultures labelled as PCE_NS_Tr2_A/B (PCE.NS.Tr2.A/B) and PCE_S_Tr2_A/B (PCE.S.Tr2.A/B) (20), grown in 50 ml per bottle, were the same cultures as indicated in step 4 in the experimental outline of previous work (20). The cultures were obtained after three spikes (see below) that were applied when PCE was completely dechlorinated into cDCE. Each spike contained 250 µM PCE and 5 mM lactate (NS and S), and 5 mM sulfate in S cultures. The actively dechlorinating cultures, PCE_NS_Tr3 and PCE_S_Tr3 (50 ml per bottle), respectively, were obtained after 5% transfer of PCE_NS_Tr2_A and PCE_S_Tr2_A. The remainders of PCE_NS_Tr2_A and PCE_S_Tr2_A were sacrificed for metagenome sequencing by Illumina (Novogene Europe, Cambridge, UK). PCE_NS_Tr2_B and PCE_S_Tr2_B cultures were used for metagenome sequencing by Nanopore separately (Novogene Europe). When 80% of the last of three spikes of 250 µM PCE was consumed, PCE_NS_Tr3 and PCE_S_Tr3 were transferred at 5 % in volume to PCE_NS_Tr4 and PCE_S_Tr4, respectively, in three replicates (100 ml per bottle). Similar to the cultures used for metagenome sequencing, three replicate cultures of PCE_NS_Tr4 and PCE_S_Tr4 were collected for metatranscriptome sequencing by Illumina (Novogene Europe) when 80% of PCE was dechlorinated into cDCE after the third spike.

### Analytical methods

PCE, TCE, and cDCE were measured by Gas chromatography and mass spectroscopy (GC-MS) installed with an Rt^®^-Q-BOND column (Retek, PA, USA) and a DSQ MS (Thermo Fisher Scientific). Hydrogen and methane were detected by compact GC (Global Analyzer Solutions, Breda, The Netherlands) with a thermal conductivity detector (GC-PDD). Organic acids, including lactate, acetate, and propionate, were measured by SHIMADZU LC2030 PLUS coupled with a Shodex SUGAR Series® SH1821 column. Sulfate was measured by Dionex™ ICS-2100 Ion Chromatography System (Thermo Scientific), and sulfide was analyzed photometrically as previously described (22).

### DNA and RNA extraction

Cultures used for DNA and RNA extraction were centrifuged at 10,000 g for 5 min, and then washed three times with 10 mM TE buffer (pH 7.0) to remove residual medium components. Washing of cultures for RNA extraction was done at 4 °C. DNA extraction was done by using the MasterPureTM Gram positive DNA purification Kit (Epicentre, WI, USA) to gain high quality and quantity DNA for metagenomic sequencing, including Illumina for short reads (PE 150, NovaSeq 6000) and Oxford Nanopore for long reads. The RNA extraction was carried out following a bead-beating procedure (23). The isolated RNA was purified using RNeasy columns (Qiagen, Venlo, The Netherlands), and residual genomic DNA was subsequently digested by DNase I (Roche, Almere, The Netherlands). The obtained RNA samples were quality-checked by agarose gel electrophoresis and sequenced using Illumina for short reads (PE 150, NovaSeq 6000).

### Metagenome data analyses

Metagenome sequence data were generated by Illumina sequencing in paired-end short reads (NovaSeq 6000) and Oxford Nanopore. The pipelines of MetaWRAP, OPERA-MS and Anvi’o were combined to process the raw data as outlined in the following (Figure S1) (24–26). One of the duplicate cultures, PCE_NS/S_Tr2_A, was sent for paired-end Illumina sequencing. Cleaning of short reads was achieved through quality check and trimming using the “read_qc” module of MetaWRAP. Both forward and reverse sequencing of PCE_NS_Tr2_A yielded 33,352,593 reads, respectively, with 33,340,280 read pairs remaining after filtering for human genomic DNA contaminants and quality control. Raw and filtered read pairs of PCE_S_Tr2_A amounted to 26,516,123 and 26,506,140, respectively. Co-assembly of the clean reads from both cultures was done using the “assembly” module of MetaWRAP. PCE_NS/S_Tr2_B cultures were used for Nanopore sequencing. The number of filtered reads of PCE_NS/S_Tr2_B was 1,637,484 and 954,603, respectively. The short-read co-assembly was then combined with the Nanopore long reads using OPERA-MS (Figure S1). Subsequently, metatranscriptome sequences (see below for processing details) were introduced to improve the quality of the hybrid assembly by OPERA-MS (Table S1). As a next step, the hybrid assembly was used for binning using the “binning” module of MetaWRAP separately with three commonly-used built-in binners: metabat2, maxbin2 and concoct. Further refining of bins was achieved using the “bin_refinement” module, and selection of bins using a cut-off at 50% completeness and 10% contamination. The abundance of the refined bins was determined using the “quant_bins” module in MetaWRAP, which uses Salmon to index the entire metagenomic assembly, and then reads from samples were mapped back to the hybrid assembly. The generated coverage estimates were used to calculate the abundance of each contig in each sample. Length-weighted average of a given bin’s contig abundances was used to calculate the bins’ abundances. Bin quality was further improved using the “reassemble_bins” module of MetaWRAP. The reassembled bins were then classified using the “classify_wf” module of GTDB-Tk (27), and phylogenomic analysis of bins was performed on the Aanovi’o platform annotated with the “annotate_bins” module of MetaWRAP using the built-in PROKKA. To assess the genomic context of target genes, flanking genes were searched by setting “grep -A 6 -B 5” to fetch the first five upstream genes and first six downstream genes using the translated “.faa” format files. The obtained gene clusters were visualized by “gggenes” (https://github.com/wilkox/gggenes).

### Phylogenetic analysis of OHR bins

Thirty-seven bins, also termed Metagenome-Assembled Genomes (MAGs) were retained after filtering using the “Bin_refinement” and “reassemble_bins” modules of MetaWRAP, with settings of > 75 % completeness and < 5 % contamination. These 37 MAGs were then phylogenomically analyzed following the Anvi’o workflow using the “--hmm-source Bacteria_71”, and the output tree file was visualized by ggtree (28), which combined the bins’ abundance, transcripts, and their genome size. Bins of potentially reductively dehalogenating bacteria were identified using the curated HMM file of RDase genes, which includes reductive dehalogenases of OHRB and tetrachloro-p-hydroquinone (TPh) reductive dehalogenase (3, 4, 29). To establish their evolutionary position, bins were taxonomically classified by “classify_wf” of GTDB (30), and bins from the same phylum or class were collected together with selected representative genomes from the respective phylum from the GTDB database (Using GTDB v202) to construct phylogenetic trees using “GToTree” (31) using 74 bacterial single-copy protein-coding genes (SCGs) for concatenated alignment. In addition, RDase entries from the Pfam database (PF13486, https://pfam.xfam.org/) were employed to retrieve all potential OHRB genomes related to bins at the phylum- or class-level. Similarly, 114 RDase protein sequences were collected from 16 bins, and multiple sequence alignments were done using the online Clustal Omega tool to construct a phylogenetic tree (32).

### Metatranscriptome data analyses

Similar to the processing of metagenome data, raw Illumina metatranscriptome sequence data was first cleaned through the removal of human genomic contaminants that may be introduced during samples preparation and extraction. Similarly, Salmon as built in the quant_bin module of MetaWRAP was also employed to calculate the abundance of bins based on transcript mapping to the respective contigs. Tuxedo packages (https://github.com/trinityrnaseq) were used for genome-guided RNA-seq analysis, including Tophat, Cufflinks, Cuffmerge and Cuffdiff (Figure S1). The hybrid assembly used for the binning (see above) was set as the template for aligning the RNA-seq data by Tophat. Cufflinks was used for assembling transcript structures from read alignments, and transcripts were counted based on Cuffdiff output, which was used for performing differential expression analysis. The abundance of standardized transcripts was taken into account to construct expression profiles of functional genes under NS and S conditions.

### Functional analyses of metabolic pathways encoded in the different bins

To evaluate the potential metabolism encoded in the 37 assembled bins, MagicLamp (https://github.com/Arkadiy-Garber/MagicLamp) was employed to search metabolic marker genes according to the established HMM files as described in “hmm-meta.txt” in the “hmms” directory (29). In our physiological experiments, we observed, alongside reductive dehalogenation, sulfate reduction as well as hydrogen and methane production, and relevant marker genes were selected. For sulfur metabolism, genes related to sulfide/sulfur/thiosulfate oxidation, sulfite/sulfate reduction, and thiosulfate disproportionation were retrieved. The initial step of sulfate reduction is the reduction of sulfate to sulfite with the formation of adenosine 5’-phosphosulfate (APS) as the intermediate, catalyzed by sulfate adenylyl-transferase (Sat) or adenylyl-sulfate kinase, and adenylyl-sulfate reductase (Apr) reducing APS to sulfite (33–35). Sulfite reductase is the critical enzyme to catalyze the reduction of sulfite to sulfide, which is the limiting step for sulfate reduction (36, 37). Two types of sulfite reductase genes were taken into consideration, including those encoding dissimilatory sulfite reductase (*dsr* genes) and anaerobic sulfite reductase (*asr* genes) (38, 39). Marker genes encoding enzymes involved in the halogen cycle were selected in addition to RDase and TPh-RDase genes, including genes encoding haloacid dehalogenase type II for organohalide breakdown, DMSO reductase type II PcrA/B for perchlorate reduction, and chlorite dismutase for chlorite reduction (40–44). Marker genes coding for hydrogenases were divided into three groups according to the metal content at active sites, i.e. [Ni-Fe]-hydrogenase, [Fe-Fe]-hydrogenase, and [Fe]-hydrogenase (45, 46). For the well-studied [Ni-Fe]-hydrogenases, eight subgroups, group 1, group 2a, group 2b, group 3a, group 3b, group 3c, group 3d, and group 4, and two subgroups of [Fe-Fe]-hydrogenases, group A and group B, were included (45–47). Metabolic genes responsible for methane production and oxidation were also searched against the assembled bins, which included *pmoA*, *pmoB*, *pmoC*, *mmoB*, and *mmoD* for methane oxidation, and *mcrA*, *mcrB*, and *mcrG* for methane production (48, 49).

## Results

### PCE dechlorination to cDCE

Stable, sediment-free PCE-dechlorinating cultures enriched from Aarhus Bay marine sediments (20) were spiked three times with PCE at 250 µM (black arrows in Figure 1B). The cultures were grown with (S) and without (NS) sulfate. The NS/S_Tr2 cultures reductively dechlorinated PCE to cDCE, reaching a final concentration of 734.7 µM and 675.1 µM respectively, after 3 spikes of 250 µM of PCE, with TCE as intermediate (Figure 1B). The resulting biomass was collected for DNA isolation for metagenome sequencing after dehalogenation of the third spike of PCE (grey arrows in Figure 1B). Similarly, the PCE_NS_Tr4 and PCE_S_Tr4 cultures reductively dechlorinated PCE to final concentrations of 636.8 µM and 528.8 µM cDCE, respectively, after 3 spikes of 250 µM of PCE. Biomass was collected when 80% of the third spike was dehalogenated to cDCE and was used for RNA extraction for metatranscriptome sequencing, thus ensuring that the OHR-associated genes were active, especially the corresponding RDase genes. The cultures from the 4th transfer without sulfate fully dechlorinated the 1st spike of PCE within 14 days, whereas the full dechlorination of the 1st spike of PCE required 30-34 days in all other sulfate-amended cultures. This is probably due to competition between sulfate and PCE as the final electron acceptor as was also observed in our previous study (20). The electron donor and carbon source, lactate, was consumed with the formation of propionate and acetate at a ratio of around 2.5 : 1 in cultures not amended with sulfate, whereas only acetate was produced in cultures where sulfate was added.

**Figure 1.**
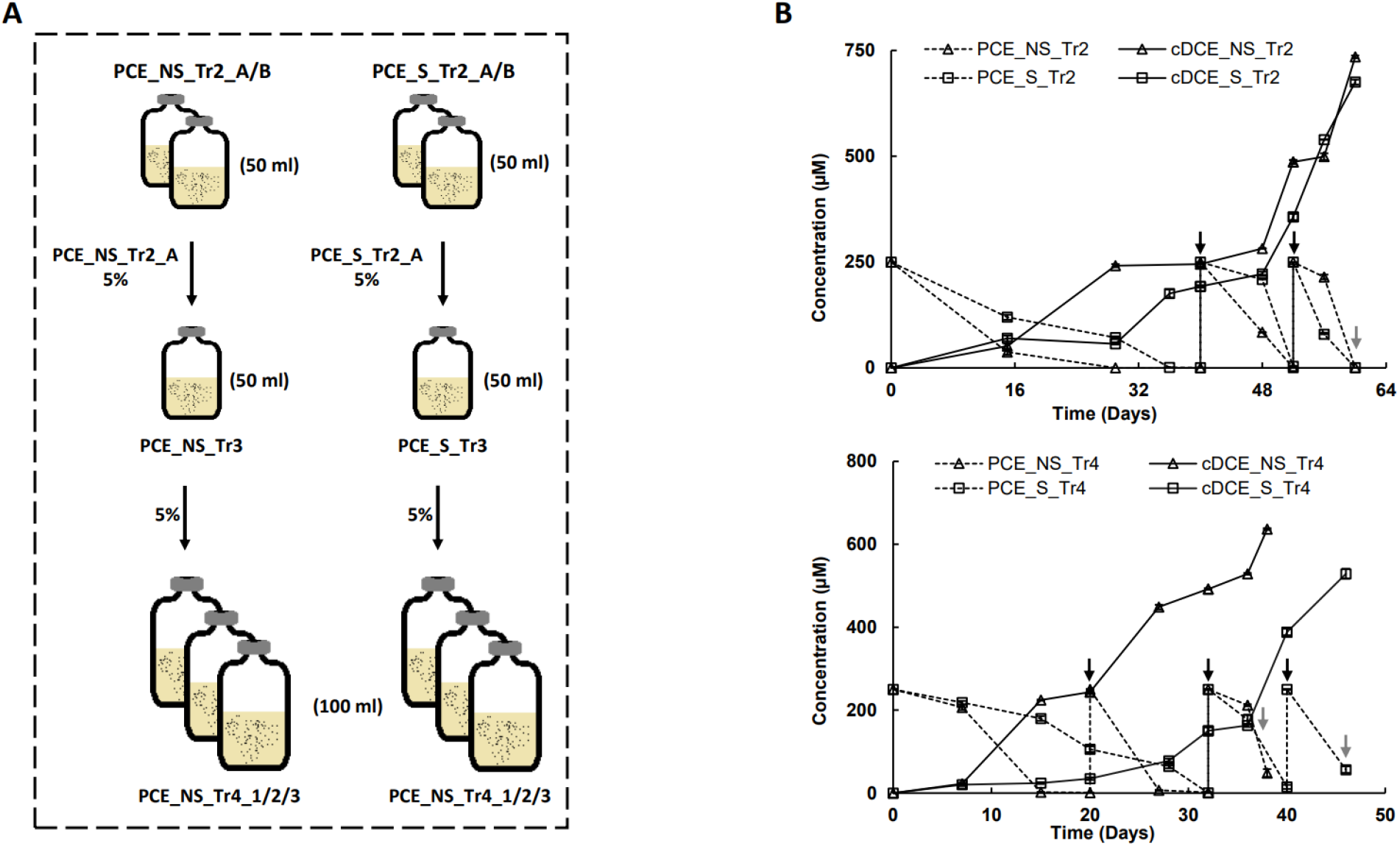
Schematic diagram (A) of PCE dechlorinating cultures (B) sampled for meta-genome and -transcriptome sequencing. PCE_NS_Tr2_A/B: duplicate PCE dechlorinating sulfate-free cultures; PCE_S_Tr2_A/B: duplicate PCE dechlorinating sulfate-amended cultures; PCE_NS/S_Tr4_1/2/3: triplicate PCE dechlorinating cultures; black arrows in B indicate PCE spikes; grey arrows represent timepoints at which cultures were harvested.

### Abundance and expression of bins in PCE dechlorinating cultures

Using the sequence data of both NS and S cultures, 37 assembled bins were retained based on completeness (>75 %) and contamination (<5%) thresholds (Table S2). Four of these bins, bin.15 (classified into genus *Desulforhopalus*), bin.22 (classified into *Desulfobacter*), bin.34 (classified into *Pseudodesulfovibrio*) and bin.5 (classified into order *Synergistales*) had 100% completeness and no contamination according to the CheckM output (25, 50). The 37 assembled bins were classified into six bacterial phyla, *Bacteroidota*, *Delongbacteria*, *Desulfobacterota*, *Firmicutes*, *Spirochaetota*, and *Synergistota*, and one archaeal phylum, *Halobacteriota*, according to the taxonomic classification following the workflow of GTDB-Tk (classify_wf) (Figure 2). The bins belonging to *Delongbacteria* and *Halobacteriota* did not encode any putative RDase genes, indicating that these organisms were probably not involved in the dechlorination of PCE.

**Figure 2.**
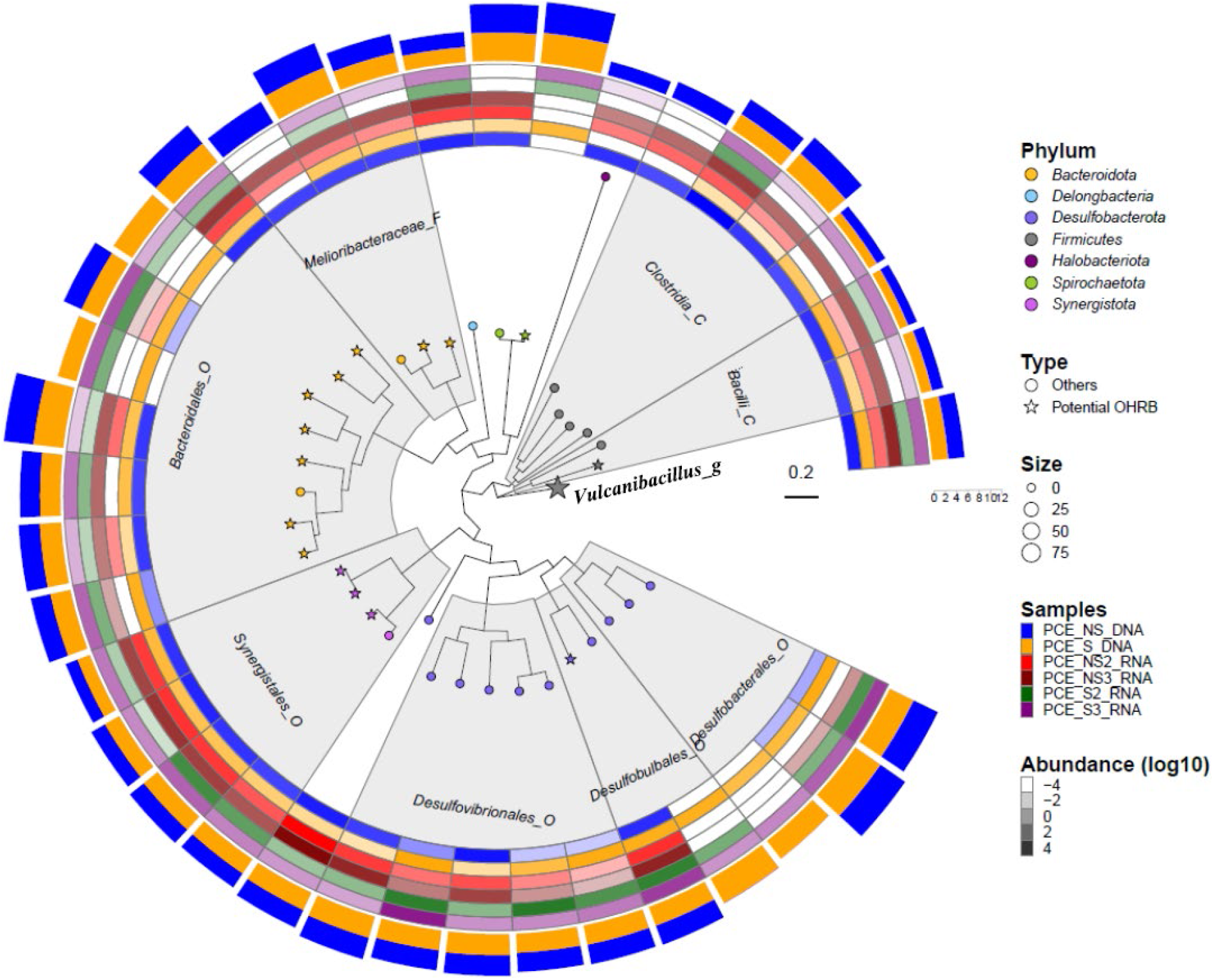
Phylogenomic tree of assembled bins constructed on the basis of 71 bacterial marker genes as implemented in Anvi’o. Tips of the tree labelled with the star symbol indicate bins containing RDase genes (“Potential OHRB”), with symbol size representing the number of RDase genes. Colors of tree tips indicate their classification at the phylum level. Areas shaded in grey indicate bins belonging to the same taxon at the level of Order (_O), Class (_C) or Family (_F). The combined heatmap in the outer circles in different colors indicate the presence of bins sourced from metagenomic (PCE_DNA) and meta-transcriptomic (PCE_RNA) data from cultures grown in the absence or presence of sulfate (NS/S). Transparency of the colors in the heatmap represents the abundance of a given bin (Unit: genome copies per million reads) in log10. The two outer circles bearing the same color pattern as the metagenomic samples indicate the presence and bar height represents the bin’s genome size as measured by the scaler (0-12 Mbp).

In the NS cultures, bin.17, classified into *Clostridiaceae*, and bin.5 classified as a member of *Synergistales* were dominant on average with 12299.6 and 3401.2 genome copies per million reads (reflecting how often a given bin is represented in the sequence data), respectively (Table S2), followed by bin.26 classified as member of *Vulcanibacillus* with 1831.6 genome copies. bin.21, classified into *Desulfuromusa* was the most highly expressed genome and accounted for 9101.4 genomic transcripts per million reads on average, followed by bin.15 classified as member of *Desulforhopalus* with 673.9 genomic transcripts per million reads. The genomes of bin.14 (classified into *Desulfomicrobium*) and bin.26 were also highly expressed with over 300 genomic transcripts per million reads.

When sulfate was present, bin.19, belonging to *Desulfoplanes*, was the most abundant and highly expressed, accounting for 6076.2 genome copies and 1145.1 genomic transcripts per million reads, followed by bin.22 with 3103.4 copies and 106.5 transcripts per million reads. bin.34, belonging to *Pseudodesulfovibrio* was the third most abundant with 3095.2 genome copies and 19.3 transcripts per million reads. bin.25, belonging to the same genus, had 333.7 transcripts as the second most highly expressed bin with 666.8 genome copies per million reads (details shown in Table S2). bin.26 belonging to the genus *Vulcanibacillus* was annotated with 97 putative RDase genes. Transcriptomics was used to check whether these genes were indeed expressed in the cultures studied. Other genomic bins containing RDase genes, bin.5, bin.10 and bin.32, belonging to *Synergistales*, bin.15 from *Desulfobacterota*, bin.28 belonging to *Melioribacteraceae*, and bin.12, bin.18, Bin.24 and bin.31 from *Bacteroidales* were present and expressed in all cultures during PCE dechlorination irrespective of the presence of sulfate, while all their correspondent RDase genes were also expressed (Figure 2; Table S4). In the sulfate amended cultures, bin.23 and bin.6, belonging to the order *Bacteroidales*, and bin.30, belonging to the *Oceanispirochaeta*, potentially contributed to the sulfate reduction, as evidenced by the detection of their genomic transcripts (Figure. 3.2, Table S2, and Table S4). In addition, the potential OHRB, bin.15 belonging to *Desulforhopalus* and bin.32 belonging to *Synergistales*, were also found to be expressed under sulfate-added condition and their correspondent RDase gene transcripts were observed with on average 65.3 (bin.15-RDase gene) and 145.5 (bin.32-RDase gene) transcripts per million reads, respectively (Table S4).

### Phylogenomic position of bins representing potential OHRB

To further assess the phylogeny of bins containing RDase genes and thus assumed to bear OHR potential, representative genomes of the different taxa were retrieved from the GTDB database to construct phylogenomic trees. Nine OHR-potential bins, bin.1, bin.12, bin.18, bin.23, bin.24, bin.28, bin.6, bin.31, and bin.35, belonging to *Bacteroidota*, were phylogenetically analyzed with representative genomes (Figure 3A). bin.1 and bin.12 belong to genus UBA12077, in which three out of six representative genomes were annotated as potential OHRB. bin.18 belongs to genus SR-FBR-E99, and no RDase was found encoded in the genomes of any of the representatives. bin.23 was classified into family VadinHA17, which included four potential OHRB representatives from genera UBA9300, JAADHC01, SLNP01 and LD21, of which UBA9300 was most closely related to bin.23. bin.31 was phylogenetically close to *Marinifilaceae*, which contained genera *Ancylomarina*, *Marinifilum*, *Ancylomarina*_A, and *Labilibaculum*. Interestingly, we found RDase genes in *Ancylomarina*, *Marinifilum*, and *Ancylomarina*_A. bin.28 and bin.35 from family *Melioribacteraceae* were close to the potential OHRB family member, JAADIR01. bin.24 was classified to genus UBA12170 that has in total two OHRB candidates across the included representatives. bin.6 was in a close relationship with genus BM520, in which one of three representatives was noted as OHRB candidate. bin.13 was classified into genus *Izemoplasma*_B in the phylum *Firmicutes*, in which all five representative genomes were potential OHRB (Figure 3B). bin.26 was affiliated with *Vulcanibacillus*, which has one isolate, *V. modesticaldus*, bearing one putative RDase gene. bin.32, belonging to the *Synergistota*, was close to *Aminiphilus*. In addition, bin.5 and bin.10 were phylogenetically close to *Aminobacterium* (Figure 3C). Within the *Desulfobacterota*, bin.15 was closest to *Desulforhopalus* that has two OHRB of the ten genome representatives (Figure 3D). bin.30 from *Spirochaeota* was classified into *Oceanispirochaeta* (Figure 3E).

**Figure 3.**
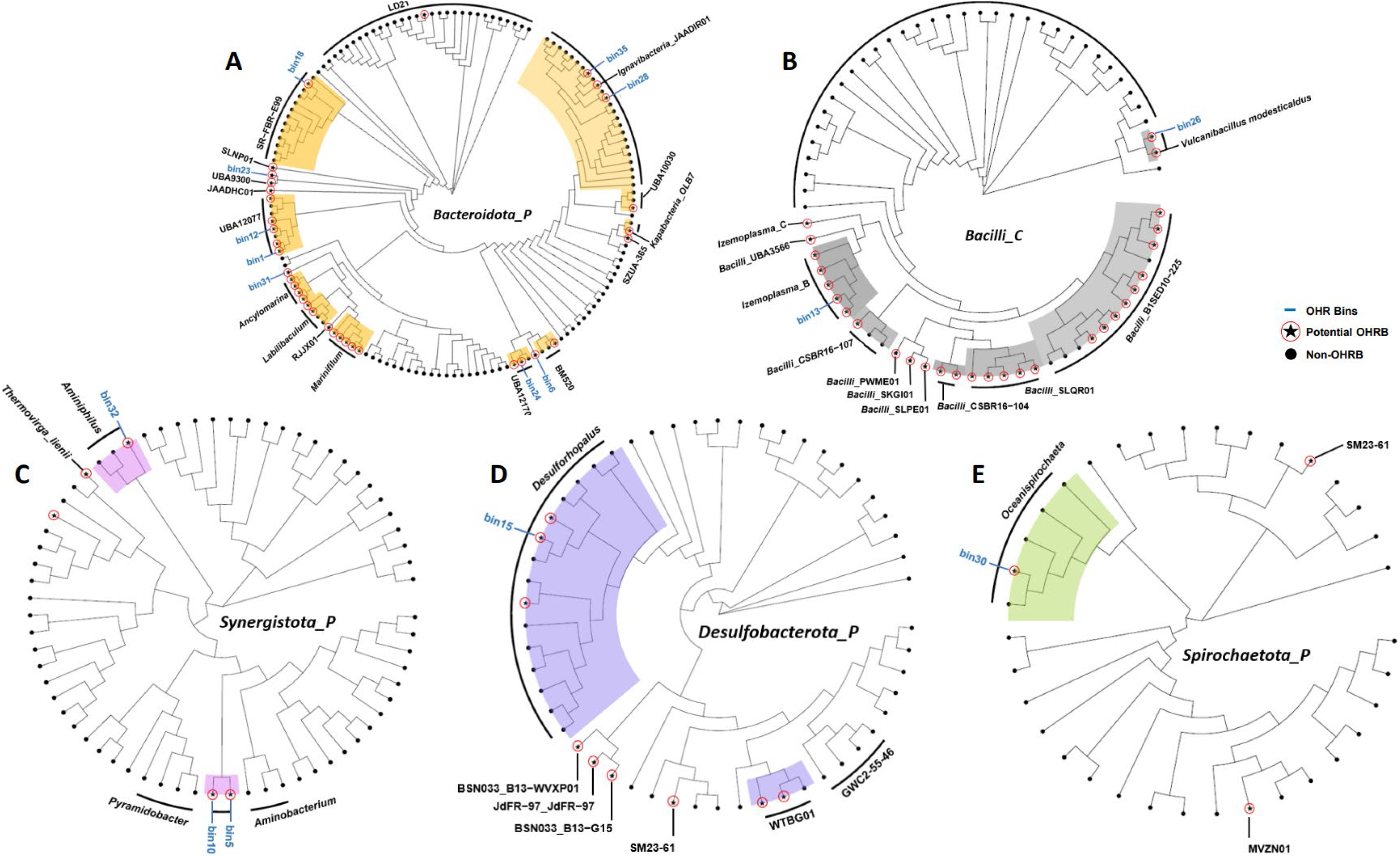
Phylogenomic trees of phyla including bins containing RDase genes. Representative genomes at phylum level: *Bacteroidota* (A), *Synergistota* (C), *Desulfobacterota* (D), and *Spirochaetota* (E) and class level: *Bacilli* (B). Red circles filled with star indicate potential OHRB, whereas black-solid dots represent non-OHRB. Colored shades indicate branches and nodes containing bins that encode RDase genes, in line with the color patterns used in Figure 1.

### Genomic survey of metabolism agreeing to physiological observations during PCE dechlorination

22 bins were found bearing metabolic genes for sulfur metabolism, of which bin.1, bin.12, bin.24, bin.28, bin.31, and bin.35 contained genes encoding both sulfate adenylyl-transferase (S_A_transferase) and adenylyl-sulfate kinase (A_S_kinase) to initiate the reduction of sulfate to sulfite with adenosine 5’-phosphosulfate (APS) as the intermediate. bin.14, bin.15, bin.16, bin.19, bin.22, bin.25, bin.27, bin.34, bin.8 and bin.9 were annotated with sulfite reductases catalyzing the further reduction of sulfite to sulfide (Figure 4D). Interestingly, there was no assembled bin bearing the complete gene set for sulfate reduction to sulfide.

**Figure 4.**
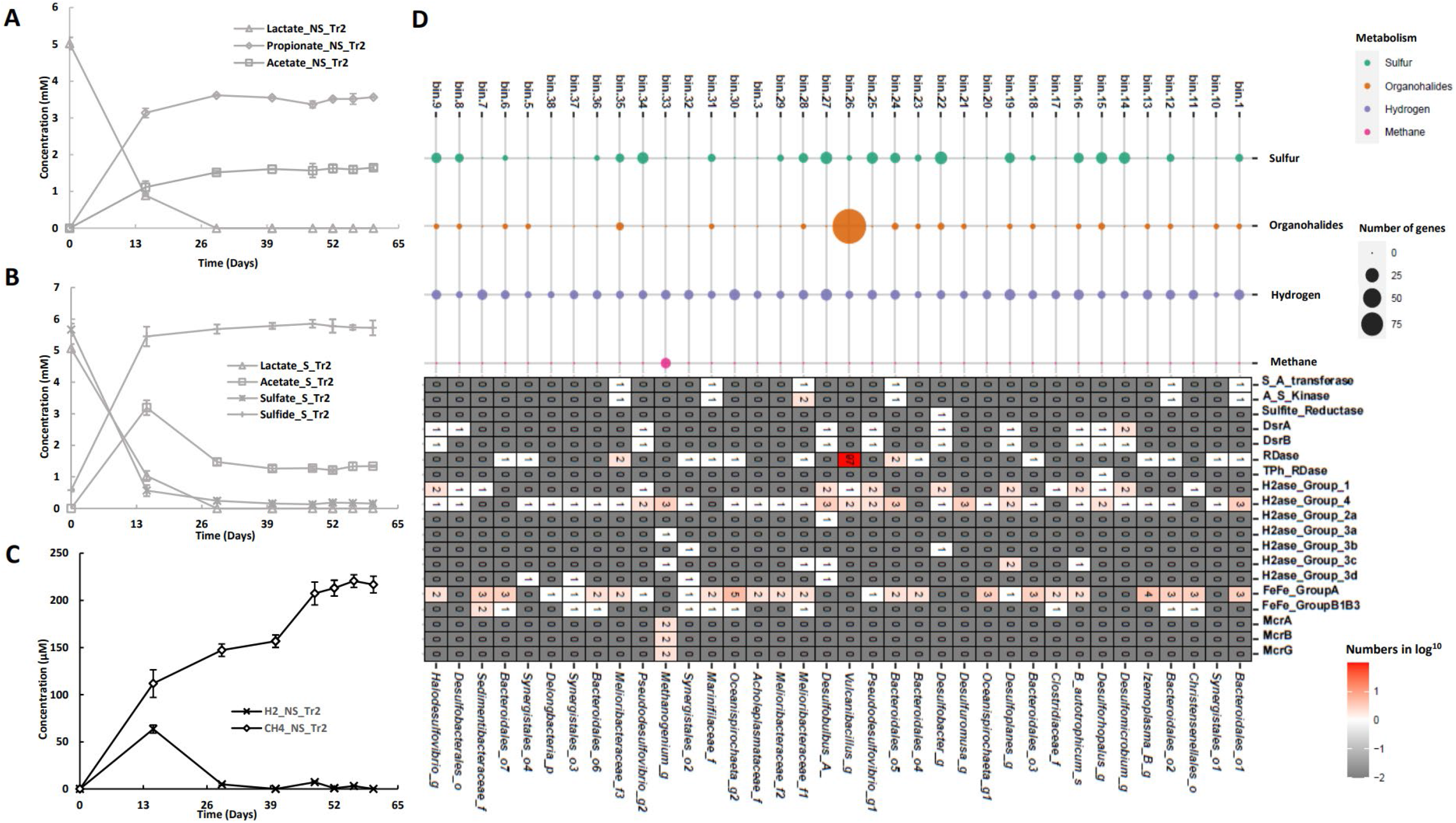
Metabolite detection and annotation of related marker genes. Utilization of lactate under sulfate-free, NS (A), and sulfate-amended, S (B) conditions. Hydrogen and methane were only detected in NS cultures (C); Marker genes involved in the cycling of sulfur, halogens, hydrogen and methane were searched against the bins and counted (D); Thereinto, specific marker genes were selected corresponding to physiological observations (D). Bars in figures A, B and C represent the standard errors of the duplicate cultures as described in Figure 1.

Metabolic marker-gene analyses revealed that there were two types of dehalogenases annotated to utilize organohalides, including RDase and TPh_RDase (Figure 4D). We identified 15 bins containing respiratory RDase genes and one TPh-RDase gene from bin.15 belonging to *Desulforhopalus*. Hydrogen was produced during lactate utilization and PCE dechlorination up to around 70 µM and consumed again after 26 days. Meanwhile, methane was also detected and accumulated to around 240 µM in the absence of added sulfate (Figure 4C). Probably, the produced hydrogen provided the electrons for reductive dehalogenation and methanogenesis under these conditions. Further genomic analysis revealed that marker genes encoding group A [Fe-Fe]-hydrogenases were abundant across the bins, for example, bin.30 and bin.13 carried five and four genes respectively, and several bins were also found to encode subgroup 1 and 4 type [Ni-Fe]-hydrogenases (Figure 4D). The higher numbers found of genes coding for [Fe-Fe]-hydrogenase suggested that they were likely the main contributors to hydrogen oxidation in accordance with previous results that [Fe-Fe]-hydrogenases were more active and had higher turnover frequency than [Ni-Fe]-hydrogenases (51). In addition, bin.23, bin.31 and bin.6 had only [Fe-Fe]-hydrogenases, whereas, bin.10, bin.14, bin.15, bin.21 and bin.26 only encoded [Ni-Fe]-hydrogenases. Genes coding for methyl-coenzyme M reductases (*mcr*) contributing to methane production were only found in bin.33, classified into *Methanogenium*, which contained two genes of each *mcrA*, *mcrB*, and *mcrG*.

### Phylogeny of RDases and their gene expression

Most RDases from bin.26 were phylogenetically closely related to each other with respect to their amino acid sequences (Table S3). RDases from other bins, except bin.35, bin.15, bin.31 and bin.24, were in a close phylogenetic relationship to each other, but distant to the RDases from bin.26 (Figure 5), suggesting a different origin. RDase gene expression profiles were analyzed to specify their possible contributions to PCE dechlorination. No RDase gene transcripts were observed for bin.35, bin.1, bin.6, bin.10, bin.12, bin.13, bin.23, bin.24, bin.28, bin.30, or bin.31. Hence, it is unlikely that those genes were involved in the dechlorination of PCE. In contrast, transcripts of several RDase genes of bin.26 were observed regardless of the presence of sulfate, including RDase3, RDase9, RDase21, RDase23, RDase43, RDase54, RDase58, RDase60, RDase61, RDase64, RDase75, RDase79 and RDase96. The RDase gene of bin.18 was transcribed only in PCE_S3 culture with 508.2 transcripts, whereas, 23 RDase genes of bin.26 were only found expressed in PCE_NS3 (Table S4). All the other RDase genes were expressed to different extents with and without sulfate. The genes from bin.15, and RDase36, RDase41, RDase42, RDase45, and RDase46 from bin.26, were expressed in all duplicate cultures (Figure 5). Notably, 84 of the 97 RDase genes from bin.26 were found expressed during PCE dechlorination, which suggests the important role of bin.26 to dehalogenate PCE in the cultures described here. In addition, most RDase genes in bin.26 were found clustered together in one assembled contig, with less than five genes interval between every two RDase genes (Table S4).

**Figure 5.**
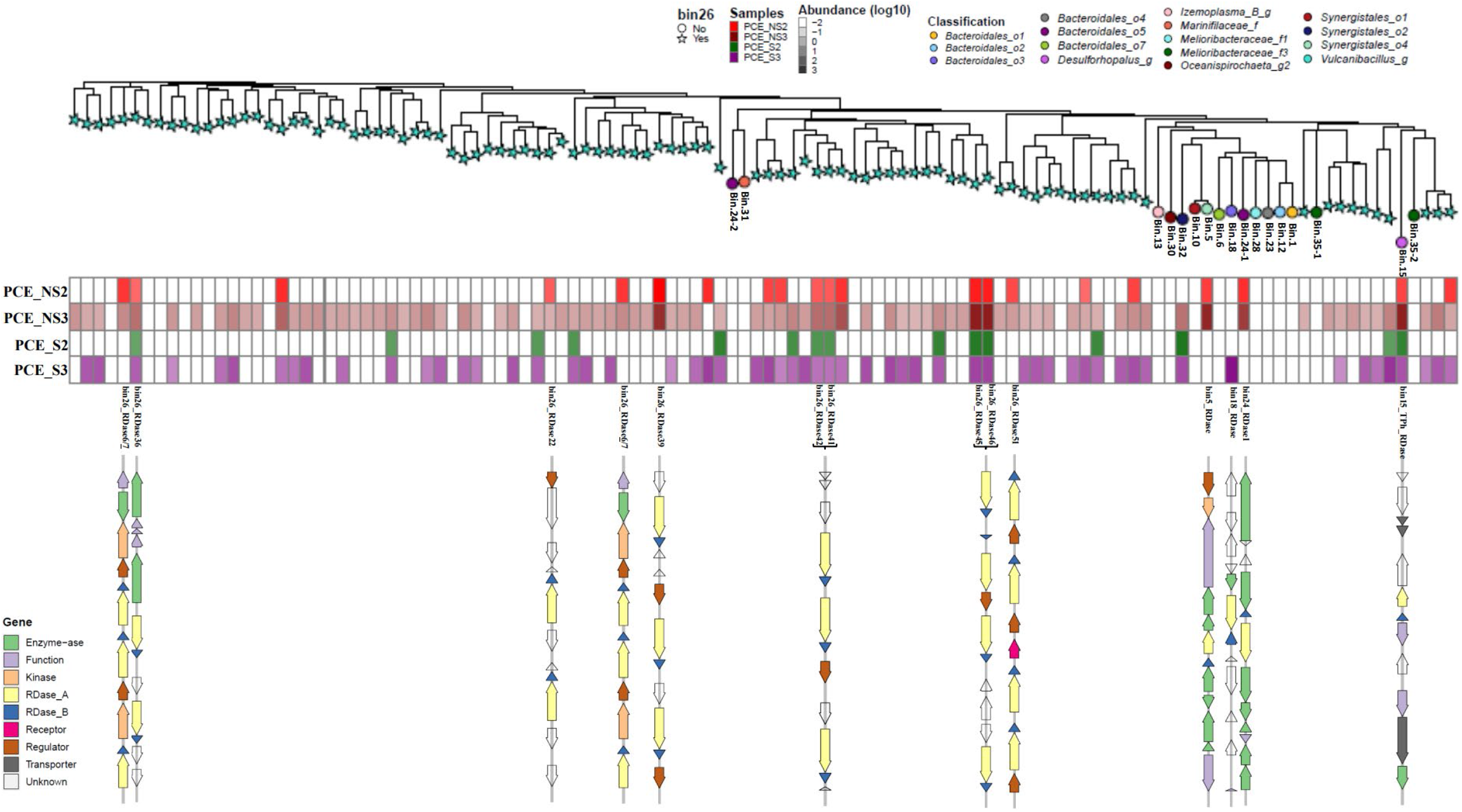
Phylogenetic tree of RDases and their gene transcripts attached with corresponding gene clusters. The protein sequences of RDases were collected and aligned via online Clustal Omega. Stars at the nodes indicate that RDases were encoded on Bin.26. RDases encoded on other bins are depicted by differently colored circles. The extended heatmap corresponding to the RDase node indicates gene transcript abundance (log10 transformed). The depicted gene clusters represent those RDase genes transcribed in both duplicates of NS and S, only in duplicates of NS, or only in S. RDase41/42, RDase45/46, and RDase6/7 are in the same cluster, respectively (Table S4).

### Diverse regulatory systems flanking RDase genes

Similar to the genes encoding RDases, the neighboring genes coding for transcriptional regulators were subjected to further analysis. CRP / FNR and TCS were the main expressed transcriptional regulators, containing 22 and 9 members, respectively (Figure 6A). CRP, the cAMP-activated global transcriptional regulator, was represented by 12 members, divided further into three groups based on sequence similarities (52). CRP-2 and CRP-3 were represented by a higher number of transcripts compared to CRP-1 (Figure 6B). Five members of GlxR (53), a CRP-like regulator, were found expressed, with GlxR1-3 exhibiting higher transcription levels compared to the rest.

**Figure 6.**
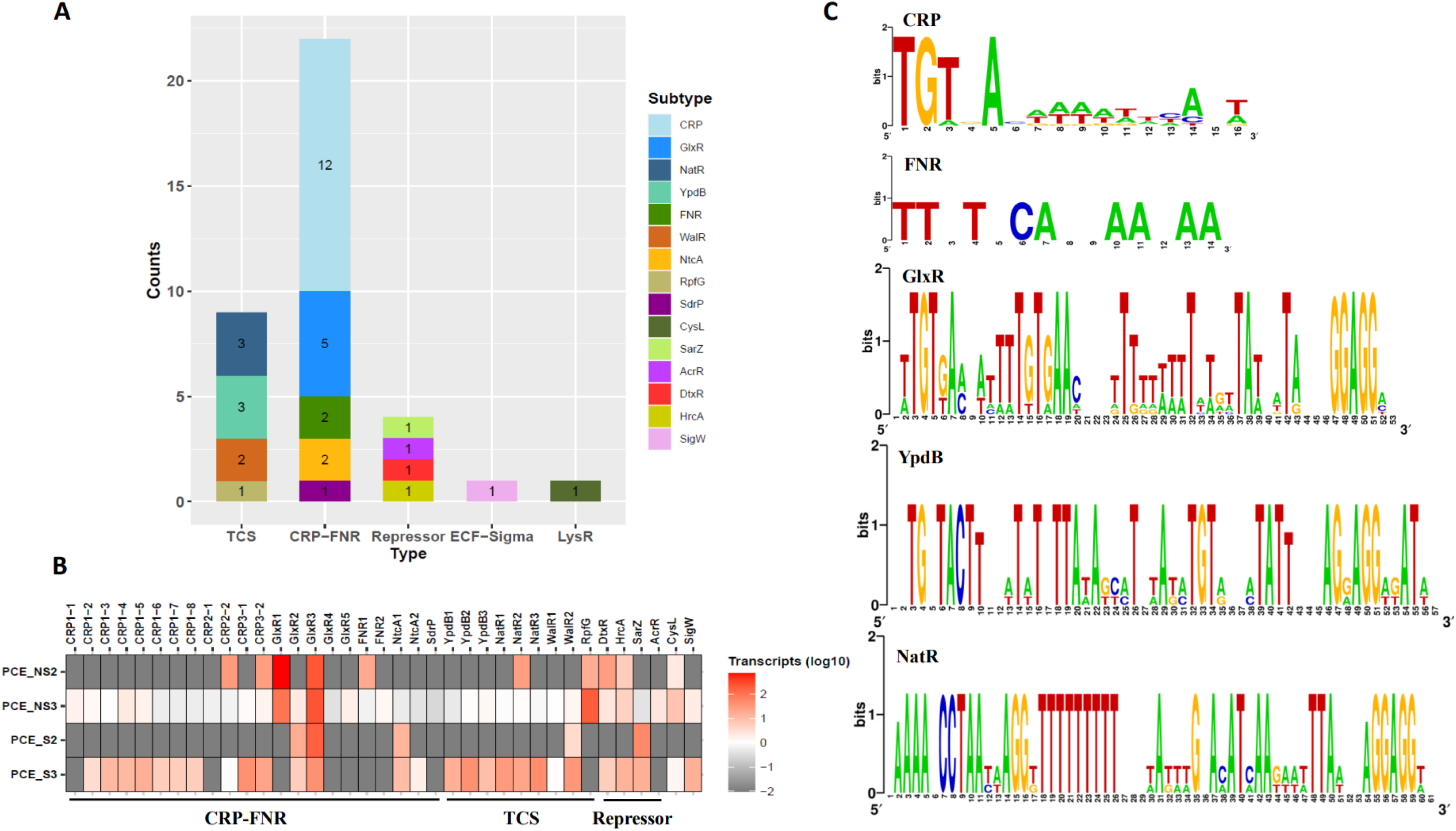
Classification and transcript abundance of expressed transcriptional regulators (A,B) and their predicted binding motifs (C). A) groups of transcriptional regulators, TCS, two-component systems; CRP-FNR, CRP (cyclic AMP receptor protein) / FNR (fumarate and nitrate reduction regulator); ECF-Sigma (ECF-σ): Extra-cytoplasmic function sigma factor; PCE_NS2 and PCE_NS3: duplicate PCE dechlorinating cultures without additional sulfate; PCE_S2 and PCE_S3: duplicate PCE dechlorinating cultures with additional sulfate; The potential binding sites were identified on the basis of binding motif in the potential promoter regions and displayed in Weblogo (https://weblogo.berkeley.edu/logo.cgi).

Nine two-component systems (TCS) were identified, of which NatR2 for regulating sodium ion extrusion (54), WalR2 for cell wall metabolism and virulence, and RpfG for signaling response, were highly expressed (Figure 6B). Interestingly, four transcriptional regulators functioning as repressors were also highly expressed, including DtxR for inhibiting Diphtheria toxin production, HrcA for preventing heat-shock induction, SarZ for attenuating virulence, and AcrR for global stresses (55–58). In addition, CysL, belonging to LysR-type regulators related with L-Cysteine biosynthesis (59), and SigW, an extra-cytoplasmic function (ECF) sigma factor responding to alkaline shock stress (60), were also induced during PCE dechlorination (Figure 6B). To better understand the regulons of these expressed transcriptional regulators, we obtained promoter sequences of RDase genes by defining intergenic regions shorter than 300bp between the RDase gene and the upstream gene (Table S5). There were 12 promoter sequences collected and each was found to contain binding sites following the consensus for CRP regulons of *Escherichia coli* (5’-TGTGACAAAATTCA*T-3’, MX000093: CRP, PRODORIC) (61) (Figure 6C). In a similar way, predicted binding sites of FNR followed the consensus 5’-TT*T*CA**AA**AA-3’ according to the two promoters of RDase genes following the consensus for known FNR regulons (MX000004: Fnr, PRODORIC). The binding sites of GlxR, YpdB, and NatR were predicted based on their RDase genes’ promoters alignment. Two promoter candidates, RDase4 and RDase70 of Bin.26, were assumed to be under the regulation of NtcA (MX000209: NtcA), of which only the promoter sequence of RDase70 was found to contain the binding site, 5’-GTATAAATATAAAC-3’.

## Discussion

Our previous work demonstrated that marine sediments from Aarhus Bay can dehalogenate a broad range of organohalides, such as PCE, under sulfate amended and sulfate free conditions (20), which for the first time provided physiological evidence of OHR for marine sediments of Aarhus Bay (17-19, 62, 63). Therefore, further pinpointing OHRB populations residing in Aarhus Bay marine sediments was the logical next step. To this end, we recruited metagenomics and -transcriptomics, yielding 37 assembled bins with high quality, of which 16 bins were predicted to represent OHRB due to their genomic annotation with RDase genes. The taxonomic classification of these OHR bins unveiled a wide distribution of RDase genes among members of *Bacteroidota*, *Snergistota*, and *Spirochaetota* in addition to the well-known phyla, such as *Firmicutes* and *Chloroflexota* (64). Gene expression analyses indicated that RDase genes of bin.26, bin.15, bin.24, bin.18, and bin.5 played the main role in PCE dechlorination, in which bin.26 was phylogenetically close to *Vulcanibacillus modesticaldus*, and bore an unprecedently high number of 97 different RDase genes, which is significantly more than the copy numbers in typical OHRB (e.g. *Dehalococcoides mccartyi* CG1 with 36 RDase genes (65)). In addition, most of the induced RDase genes were clustered with genes encoding diverse regulatory systems suggesting their strong flexibility adapting to environmental changes or the availability of different organohalides.

### Wide distribution of RDases beyond well-characterized OHRB

Most of the OHRB were isolated from organohalide-contaminated areas, such as soils, rivers and lakes (66–68). They belong to the phyla *Chloroflexota*, *Firmicutes*, *Desulfobacterota*, and *Proteobacteria* (64, 69). Although pristine marine sediments have been shown to contain OHRB such as *Dehalococcoides* (70), fewer OHRB were isolated from pristine marine environments, which might be due to the limited knowledge of their metabolic potential and limited methods for isolation. The employed integration of metagenomics and metatranscriptomics in this study allowed for the assembly of 15 OHR bins with high quality, some of which belonging to *Bacteroidota*, *Spirochaetota*, and *Synergistota* phyla that have never been reported to catalyze reductive dehalogenation. The average nucleotide identity (ANI) of these OHR bins was below 95% in all cases after running the “classify_wf” of GTDB, indicating they are novel genomes. Our further exploration together with representative genomes of these phyla revealed that 112 genomes of *Bacteroidota*, 15 genomes of *Spirochaetota*, and seven genomes of *Synergistota* are bearing RDase genes, with none of the corresponding isolates being physiologically characterized to perform OHR. To this end, it is noteworthy that our study using genome-resolved strategies discovered that the RDase genes from bin.18, classified into *Bacteroidota*, and from bin.5 and bin.32, classified into *Synergistota*, were expressed in PCE dechlorinating cultures (Table S2), and their genome abundances and genomic transcripts were in high numbers, suggesting that OHR potential exists in these phyla. The RDase genes in the other bins from the above phyla were not found expressed in our data, indicating that they were probably not involved in the respiration of PCE in our experiments. We found 32 representative genomes from the class *Bacilli* that are annotated with RDase genes. Bin.26 is closely related to *Vulcanibacillus* which has one isolate from a moderately hydrothermal vent containing one RDase gene (71). In contrast, bin.26 was predicted to contain 97 RDase genes, a number that is higher than the maximum number of 36 currently reported for members of *Dehalococcoides* (65, 72, 73). *Desulfobacterota* is a reclassified phylum from Deltaproteobacteria and well known for catalyzing sulfate reduction (69). Eighty-nine of the 939 representative genomes in the *Desulfobacterota* were found to carry RDase genes, similar to previously described strains of *Desulfoluna*, *Desulfuromusa*, and *Desulfovibrio* (21, 74). Bin.21, classified into *Desulfuromusa*, had no annotated genes involved in OHR or sulfate reduction and had a high abundance when sulfate was absent as was also found in the preceding study (20). Bin.21 only has three group 4 [Ni-Fe]-hydrogenase genes, and could thus act as a potential hydrogen producer in our cultures. Noticeably, the higher numbers found of genes coding for [Fe-Fe]-hydrogenase suggested that they were likely the main contributors to hydrogen production in accordance with previous results that [Fe-Fe]-hydrogenases were more active and had higher turnover frequency than [Ni-Fe]-hydrogenases (51). Bin.15 was most closely affiliated with the genus *Desulforhopalus*, of which one putative OHRB is *D. singaporensis* (75). Noticeably, the deduced RDase from bin.15 does not belong to the canonical RDases associated with OHR, such as PceA from *Dehalococcoides* (76, 77), and was more similar in sequence and size to TPh-RDase from *Sphingobium chlorophenolicum* (previously named as *Flavobacterium* sp.) that can dehalogenate tetrachlorohydroquinone thiolytically (3, 4). Of interest, the TPh-RDase from bin.15 was not reported for respiratory dehalogenation to conserve energy, however, the transcript data revealed TPh-RDase bears potential biodegrading activity of PCE in the cultures studied here. Moreover, the genomic abundance and transcript of bin.15 indicated its participation in PCE dechlorination. Thus, further metabolic analysis of bin.15 is essential. Altogether, marine sediments of Aarhus Bay are home to several new OHRB candidates with diverse RDase genes.

### RDase gene clusters

Recently, an extensive genomic survey found that some RDase gene clusters lack genes encoding anchor protein B, or the N-terminus of RDase bears transmembrane domains, which could leave the RDase functioning in the cytoplasm or the C-terminus spanning towards the outer face of the membrane (78, 79). In our study, we also observed some OHR bins without putative RdhB encoding genes, including bin.12, bin.1, bin.28, bin.32 and bin.6. Of these, transcripts of the RDase gene in bin.32 indicated the possible cytoplasmic dehalogenation that awaits further experimental confirmation. In most cases, RDase genes were accompanied by diverse functional gene sets, such as cobamide cofactor biosynthesis pathway genes in *Sulfurospirillum* strains and molecular chaperones in *Desulfitobacterium* (80–82). Similarly, some RDase gene clusters found in the present study contained chaperone genes vicinal to genes encoding RDase3, RDase4, RDase35, RDase93 of bin.26 that could protect the RDase activity in a manner of maintaining structural integrity when exposed to harsh environments. Besides, there were several genes encoding electron transport complexes accompanying RDase genes, including RDase in bin.13, bin.31, RDase2 in bin.24 and bin.35, and RDase4, RDase67, RDase69, RDase73, RDase82, and RDase84 in bin.26, which could form new electron transport chains to promote OHR in addition to the previously identified ones (7, 83).

Furthermore, RDase genes were frequently flanked by genes coding for transcriptional regulators, which might timely and accurately regulate the expression of vicinal RDase genes in response to the added organohalides. Three regulatory systems were previously characterized, including those of the CRP-FNR family, MarR and TCS (5, 6, 81, 82, 84). Expressed transcriptional regulators classified into CRP/FNR systems accounted for the largest numbers in our study, whereas, GlxR, as a new regulator, showed different regulons inferred from the promoter alignment as well as YpdB and NatR from the class of TCS. In addition, four negative regulators were found that could follow the modulation reported for MarR-type regulators (6), but their binding sites were unclear and need further demonstration. Noticeably, ECF sigma factor, SigW, was also found transcribed during PCE dechlorination, which was rarely reported to regulate dehalogenation, and its DNA binding motif seems different from that in *Bacillus subtilis* (MX000079: SigW). Taken together, the newly-discovered regulatory systems diversified the known modes of regulation of dehalogenation that could be the result of adaptation to challenging environments, and further molecular identification would substantially build up our understanding on reductive dehalogenation and its regulation in natural systems.

## Supporting information

Caption for supplementary tables is described in the manuscript

## Data availability

The nucleotide sequence data has been deposited in the European Bioinformatics Institute under accession number PRJEB61282.

## Author contributions

C.Z., T.B., S.A., and H.S. conceived and coordinated the study. C.Z. wrote the primary manuscript and performed all experiments and data analysis. T.B., S.A., and H.S. gave detailed revision and C.Z. completed the manuscript.

## Acknowledgements

This study was supported by the Dutch Research Council through the UNLOCK project (NRGWI.obrug.2018.005), as well as Wageningen University through its Innovation Program Microbiology. We thank Felix Homa and Victor de Jager (Laboratory of Microbiology, Wageningen University) for providing help for the metagenomic and meta-transcriptomic analyses. We furthermore would like to thank Sjoerd Engelbertink for GC-MS measurements. We acknowledge the China Scholarship Council (CSC) for the support to Chen Zhang (File No. 201807720048).

## Supplementary information

**Figure S1.**
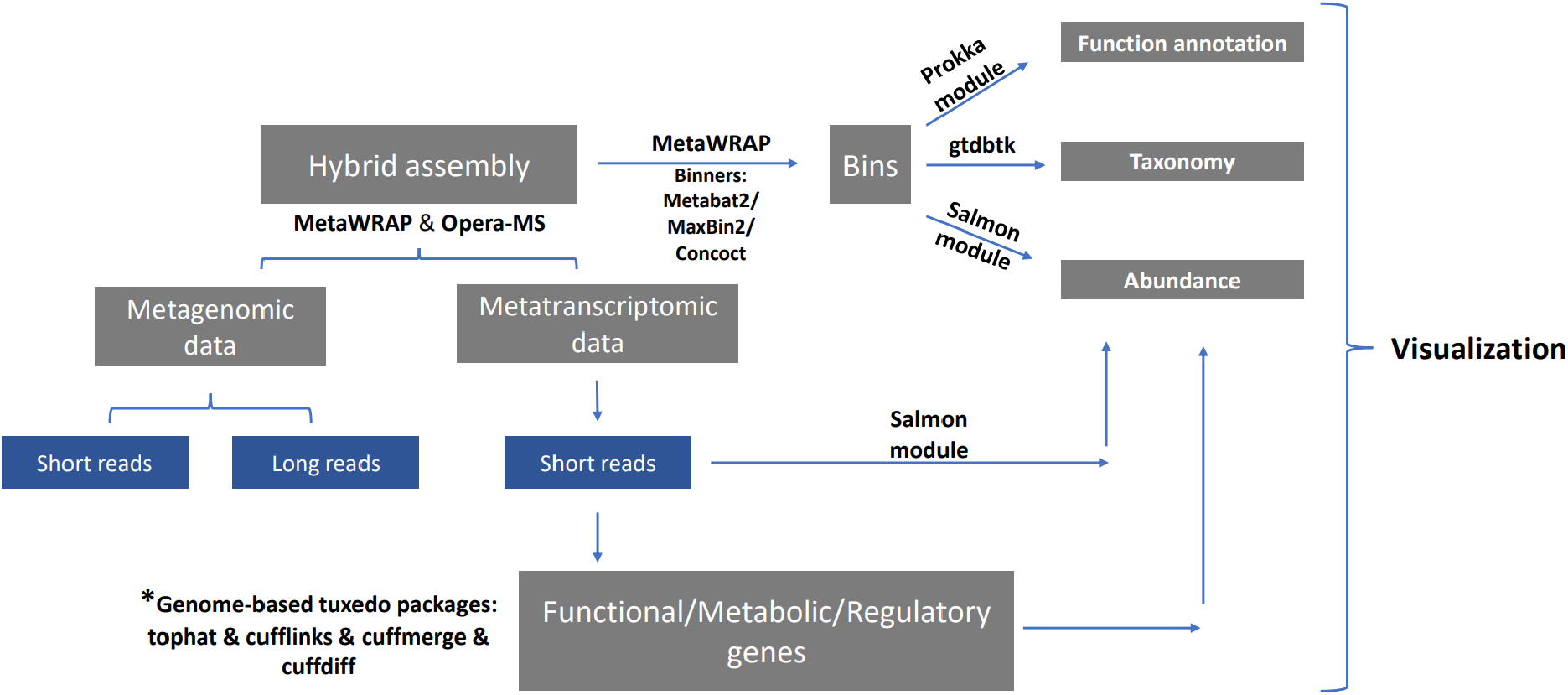
Outline of metagenomic and meta-transcriptomic sequence processing and analyses. * Analyses of metatranscriptomic data was performed by tuxedo packages that include tophat (aligning reads), cufflinks (assembling transcripts), cuffmerge (merging transcripts) and cuffdiff (identifying differentially-expressed transcripts) (85). Arrows in blue indicate stepwise analyses. Visualization of output data was achieved using R and the required packages, including ggplot2, ggtree, and gggenes.

## Supplementary tables

**Table S1** Metagenomic assembly, using short and long read DNA sequences as well as short read metatranscriptome sequences

**Table S2** Metagenome Assembled Genomes (MAGs) with abundance of genomic copies and transcripts in per million reads unit

**Table S3** Protein sequences of RDases among bins

**Table S4** Transcriptional profile of RDase genes in OHR bins

**Table S5** Predicted promoter sequences of RDase genes

## References

1. Adrian L, Löffler FE. 2016. Organohalide-respiring bacteria, vol 85. Springer.

2. Mohn WW, Tiedje J. 1992. Microbial reductive dehalogenation. Microbiol Mol Biol Rev 56:482–507.

3. Xun L, Topp E, Orser C. 1992. Glutathione is the reducing agent for the reductive dehalogenation of tetrachloro-p-hydroquinone by extracts from a *Flavobacterium* sp. Biochem Biophys Res Commun 182:361–366.

4. Xun L, Topp E, Orser C. 1992. Purification and characterization of a tetrachloro-p-hydroquinone reductive dehalogenase from a *Flavobacterium* sp. J Bacteriol 174:8003–8007.

5. Gabor K, Verissimo CS, Cyran BC, Ter Horst P, Meijer NP, Smidt H, De Vos WM, Van der Oost J. 2006. Characterization of CprK1, a CRP/FNR-type transcriptional regulator of halorespiration from *Desulfitobacterium hafniense*. J Bacteriol 188:2604–13.

6. Wagner A, Segler L, Kleinsteuber S, Sawers G, Smidt H, Lechner U. 2013. Regulation of reductive dehalogenase gene transcription in *Dehalococcoides mccartyi*. Philos Trans R Soc Lond B Biol Sci 368:20120317.

7. Wang S, Qiu L, Liu X, Xu G, Siegert M, Lu Q, Juneau P, Yu L, Liang D, He Z, Qiu R. 2018. Electron transport chains in organohalide-respiring bacteria and bioremediation implications. Biotechnol Adv 36:1194–1206.

8. Fincker M, Spormann AM. 2017. Biochemistry of catabolic reductive dehalogenation. Annu Rev Biochem 86:357–386.

9. Smidt H, De Vos WM. 2004. Anaerobic microbial dehalogenation. Annu Rev Microbiol 58:43–73.

10. Peng P, Lu Y, Bosma TNP, Nijenhuis I, Nijsse B, Shetty SA, Ruecker A, Umanets A, Ramiro-Garcia J, Kappler A, Sipkema D, Smidt H, Atashgahi S. 2020. Metagenomic- and cultivation-based exploration of anaerobic chloroform biotransformation in hypersaline sediments as natural source of chloromethanes. Microorganisms 8:665.

11. Atashgahi S, Haggblom MM, Smidt H. 2018. Organohalide respiration in pristine environments: implications for the natural halogen cycle. Environ Microbiol 20:934–948.

12. Gribble GW. 2015. A recent survey of naturally occurring organohalogen compounds. Environ Chem 12:396–405.

13. Smits TH, Devenoges C, Szynalski K, Maillard J, Holliger C. 2004. Development of a real-time PCR method for quantification of the three genera *Dehalobacter*, *Dehalococcoides*, and *Desulfitobacterium* in microbial communities. J Microbiol Methods 57:369–78.

14. Lu Y, Atashgahi S, Hug LA, Smidt H. 2017. Primers that target functional genes of organohalide-respiring bacteria. Hydrocarbon and Lipid Microbiology Protocols: Primers:177–205.

15. Bokulich NA, Kaehler BD, Rideout JR, Dillon M, Bolyen E, Knight R, Huttley GA, Gregory Caporaso J. 2018. Optimizing taxonomic classification of marker-gene amplicon sequences with QIIME 2’s q2-feature-classifier plugin. Microbiome 6:1–17.

16. Spang A, Stairs CW, Dombrowski N, Eme L, Lombard J, Caceres EF, Greening C, Baker BJ, Ettema TJG. 2019. Proposal of the reverse flow model for the origin of the eukaryotic cell based on comparative analyses of *Asgard* archaeal metabolism. Nat Microbiol 4:1138–1148.

17. Wasmund K, Cooper M, Schreiber L, Lloyd KG, Baker BJ, Petersen DG, Jorgensen BB, Stepanauskas R, Reinhardt R, Schramm A, Loy A, Adrian L. 2016. Single-cell genome and group-specific *dsrAB* sequencing implicate marine members of the class *Dehalococcoidia* (Phylum *Chloroflexi*) in sulfur cycling. mBio 7.

18. Zinke LA, Mullis MM, Bird JT, Marshall IPG, Jorgensen BB, Lloyd KG, Amend JP, Kiel Reese B. 2017. Thriving or surviving? Evaluating active microbial guilds in Baltic Sea sediment. Environ Microbiol Rep 9:528–536.

19. Jochum LM, Schreiber L, Marshall IPG, Jorgensen BB, Schramm A, Kjeldsen KU. 2018. Single-cell genomics reveals a diverse metabolic potential of uncultivated *Desulfatiglans*-related *Deltaproteobacteria* widely distributed in marine sediment. Front Microbiol 9:2038.

20. Zhang C, Atashgahi S, Bosma TNP, Peng P, Smidt H. 2022. Organohalide respiration potential in marine sediments from Aarhus Bay. FEMS Microbiol Ecol 98:fiac073.

21. Liu J, Haggblom MM. 2018. Genome-guided identification of organohalide-respiring *Deltaproteobacteria* from the marine environment. mBio 9:e02471–18.

22. Cline JD. 1969. Spectrophotometric determination of hydrogen sulfide in natural waters. Limnol Oceanogr 14:454–458.

23. Egert M, De Graaf AA, Maathuis A, De Waard P, Plugge CM, Smidt H, Deutz NE, Dijkema C, De Vos WM, Venema K. 2007. Identification of glucose-fermenting bacteria present in an in vitro model of the human intestine by RNA-stable isotope probing. FEMS Microbiol Ecol 60:126–135.

24. Eren AM, Kiefl E, Shaiber A, Veseli I, Miller SE, Schechter MS, Fink I, Pan JN, Yousef M, Fogarty EC. 2021. Community-led, integrated, reproducible multi-omics with anvi’o. Nat Microbiol 6:3–6.

25. Uritskiy GV, DiRuggiero J, Taylor J. 2018. MetaWRAP—a flexible pipeline for genome-resolved metagenomic data analysis. Microbiome 6:1–13.

26. Bertrand D, Shaw J, Kalathiyappan M, Ng AHQ, Kumar MS, Li C, Dvornicic M, Soldo JP, Koh JY, Tong C. 2019. Hybrid metagenomic assembly enables high-resolution analysis of resistance determinants and mobile elements in human microbiomes. Nat Biotechnol 37:937–944.

27. Rinke C, Chuvochina M, Mussig AJ, Chaumeil P-A, Davín AA, Waite DW, Whitman WB, Parks DH, Hugenholtz P. 2021. A standardized archaeal taxonomy for the Genome Taxonomy Database. Nat Microbiol 6:946–959.

28. Yu G, Smith DK, Zhu H, Guan Y, Lam TTY. 2017. ggtree: an R package for visualization and annotation of phylogenetic trees with their covariates and other associated data. Methods Ecol Evol 8:28–36.

29. Garber A, Ramirez G, Merino N, Pavia M, McAllister S. 2020. MagicLamp: toolkit for annotation of ‘omics datasets using curated HMM sets. GitHub Repository Available online at https://githubcom/Arkadiy-Garber/MagicLamp (accessed January, 2022).

30. Chaumeil P-A, Mussig AJ, Hugenholtz P, Parks DH. 2020. GTDB-Tk: a toolkit to classify genomes with the Genome Taxonomy Database. Bioinformatics 36:1925–1927.

31. Lee MD. 2019. GToTree: a user-friendly workflow for phylogenomics. Bioinformatics 35:4162–4164.

32. Sievers F, Wilm A, Dineen D, Gibson TJ, Karplus K, Li W, Lopez R, McWilliam H, Remmert M, Söding J. 2011. Fast, scalable generation of high-quality protein multiple sequence alignments using Clustal Omega. Mol Syst Biol 7:539.

33. Leloup J, Fossing H, Kohls K, Holmkvist L, Borowski C, Jorgensen BB. 2009. Sulfate-reducing bacteria in marine sediment (Aarhus Bay, Denmark): abundance and diversity related to geochemical zonation. Environ Microbiol 11:1278–91.

34. Muyzer G, Stams AJ. 2008. The ecology and biotechnology of sulphate-reducing bacteria. Nat Rev Microbiol 6:441–54.

35. Anantharaman K, Hausmann B, Jungbluth SP, Kantor RS, Lavy A, Warren LA, Rappé MS, Pester M, Loy A, Thomas BC. 2018. Expanded diversity of microbial groups that shape the dissimilatory sulfur cycle. ISME J 12:1715–1728.

36. Leavitt WD, Bradley AS, Santos AA, Pereira IA, Johnston DT. 2015. Sulfur isotope effects of dissimilatory sulfite reductase. Front Microbiol 6:1392.

37. Bradley A, Leavitt W, Johnston D. 2011. Revisiting the dissimilatory sulfate reduction pathway. Geobiology 9:446–457.

38. Hallenbeck PC, Clark MA, Barrett EL. 1989. Characterization of anaerobic sulfite reduction by *Salmonella typhimurium* and purification of the anaerobically induced sulfite reductase. J Bacteriol 171:3008–3015.

39. Peck Jr H. 1961. Enzymatic basis for assimilatory and dissimilatory sulfate reduction. J Bacteriol 82:933–939.

40. Hisano T, Hata Y, Fujii T, Liu J-Q, Kurihara T, Esaki N, Soda K. 1996. Crystal structure of L-2-Haloacid dehalogenase from *Pseudomonas* sp. YL: an α/β hydrolase structure that is different from the α/β hydrolase fold. J Biol Chem 271:20322–20330.

41. Koonin EV, Tatusov RL. 1994. Computer analysis of bacterial haloacid dehalogenases defines a large superfamily of hydrolases with diverse specificity: application of an iterative approach to database search. J Mol Biol 244:125–132.

42. Peng P, Zheng Y, Koehorst J, Schaap P, Stams A, Smidt H, Atashgahi S. 2017. Concurrent haloalkanoate degradation and chlorate reduction by *Pseudomonas chloritidismutans* AW-1T. Appl Environ Microbiol 83:e00325–17.

43. Bender KS, Shang C, Chakraborty R, Belchik SM, Coates JD, Achenbach LA. 2005. Identification, characterization, and classification of genes encoding perchlorate reductase. J Bacteriol 187:5090–5096.

44. Coates JD, Achenbach LA. 2004. Microbial perchlorate reduction: rocket-fuelled metabolism. Nat Rev Microbiol 2:569–580.

45. Piché-Choquette S, Constant P. 2019. Molecular hydrogen, a neglected key driver of soil biogeochemical processes. Appl Environ Microbiol 85:e02418–18.

46. Lubitz W, Ogata H, Rudiger O, Reijerse E. 2014. Hydrogenases. Chem Rev 114:4081–4148.

47. Schoelmerich MC, Muller V. 2019. Energy-converting hydrogenases: the link between H_2_ metabolism and energy conservation. Cell Mol Life Sci doi:10.1007/s00018-019-03329-5.

48. Hakemian AS, Rosenzweig AC. 2007. The biochemistry of methane oxidation. Annu Rev Biochem 76:223–241.

49. Evans PN, Boyd JA, Leu AO, Woodcroft BJ, Parks DH, Hugenholtz P, Tyson GW. 2019. An evolving view of methane metabolism in the Archaea. Nat Rev Microbiol 17:219–232.

50. Parks DH, Imelfort M, Skennerton CT, Hugenholtz P, Tyson GW. 2015. CheckM: assessing the quality of microbial genomes recovered from isolates, single cells, and metagenomes. Genome Res 25:1043–1055.

51. Madden C, Vaughn MD, Díez-Pérez I, Brown KA, King PW, Gust D, Moore AL, Moore TA. 2012. Catalytic turnover of [FeFe]-hydrogenase based on single-molecule imaging. J Am Chem Soc 134:1577–1582.

52. Chandler MS. 1992. The gene encoding cAMP receptor protein is required for competence development in *Haemophilus influenzae* Rd. Proc Natl Acad Sci USA 89:1626–1630.

53. Kim H-J, Kim T-H, Kim Y, Lee H-S. 2004. Identification and characterization of *glxR*, a gene involved in regulation of glyoxylate bypass in *Corynebacterium glutamicum*. J Bacteriol 186:3453–3460.

54. Ogura M, Tsukahara K, Hayashi K, Tanaka T. 2007. The *Bacillus subtilis* NatK–NatR two-component system regulates expression of the *natAB* operon encoding an ABC transporter for sodium ion extrusion. Microbiology 153:667–675.

55. Kaito C, Morishita D, Matsumoto Y, Kurokawa K, Sekimizu K. 2006. Novel DNA binding protein SarZ contributes to virulence in *Staphylococcus aureus*. Mol Microbiol 62:1601–1617.

56. Narberhaus F. 1999. Negative regulation of bacterial heat shock genes. Mol Microbiol 31:1–8.

57. Pohl E, Holmes RK, Hol WG. 1999. Crystal structure of a cobalt-activated diphtheria toxin repressor-DNA complex reveals a metal-binding SH3-like domain. J Mol Biol 292:653–667.

58. Ma D, Alberti M, Lynch C, Nikaido H, Hearst JE. 1996. The local repressor AcrR plays a modulating role in the regulation of acrAB genes of *Escherichia coli* by global stress signals. Mol Microbiol 19:101–112.

59. Guillouard I, Auger S, Hullo M-Fo, Chetouani F, Danchin A, Martin-Verstraete I. 2002. Identification of *Bacillus subtilis* CysL, a regulator of the cysJI operon, which encodes sulfite reductase. J Bacteriol 184:4681–4689.

60. Wiegert T, Homuth G, Versteeg S, Schumann W. 2001. Alkaline shock induces the Bacillus subtilis σ^W^ regulon. Mol Microbiol 41:59–71.

61. Dudek C-A, Jahn D. 2022. PRODORIC: state-of-the-art database of prokaryotic gene regulation. Nucleic Acids Res 50:D295–D302.

62. Wasmund K, Algora C, Muller J, Kruger M, Lloyd KG, Reinhardt R, Adrian L. 2015. Development and application of primers for the class *Dehalococcoidia* (phylum *Chloroflexi*) enables deep insights into diversity and stratification of subgroups in the marine subsurface. Environ Microbiol 17:3540–56.

63. Wasmund K, Schreiber L, Lloyd KG, Petersen DG, Schramm A, Stepanauskas R, Jorgensen BB, Adrian L. 2014. Genome sequencing of a single cell of the widely distributed marine subsurface *Dehalococcoidia*, phylum *Chloroflexi*. ISME J 8:383–97.

64. Atashgahi S, Lu Y, Smidt H. 2016. Overview of known organohalide-respiring bacteria—phylogenetic diversity and environmental distribution, p 63–105. In Adrian L, Löffler FE (ed), Organohalide-Respiring Bacteria doi:10.1007/978-3-662-49875-0_5. Springer Berlin Heidelberg, Berlin, Heidelberg.

65. Molenda O, Puentes Jacome LA, Cao X, Nesbo CL, Tang S, Morson N, Patron J, Lomheim L, Wishart DS, Edwards EA. 2020. Insights into origins and function of the unexplored majority of the reductive dehalogenase gene family as a result of genome assembly and ortholog group classification. Environ Sci Process Impacts doi:10.1039/c9em00605b.

66. Wiegel J, Wu Q. 2000. Microbial reductive dehalogenation of polychlorinated biphenyls. FEMS Microbiol Ecol 32:1–15.

67. Lee LK, He J. 2010. Reductive debromination of polybrominated diphenyl ethers by anaerobic bacteria from soils and sediments. Appl Environ Microbiol 76:794–802.

68. Chambon JC, Bjerg PL, Scheutz C, Bælum J, Jakobsen R, Binning PJ. 2013. Review of reactive kinetic models describing reductive dechlorination of chlorinated ethenes in soil and groundwater. Biotechnol Bioeng 110:1–23.

69. Waite DW, Chuvochina M, Pelikan C, Parks DH, Yilmaz P, Wagner M, Loy A, Naganuma T, Nakai R, Whitman WB, Hahn MW, Kuever J, Hugenholtz P. 2020. Proposal to reclassify the proteobacterial classes *Deltaproteobacteria* and *Oligoflexia*, and the phylum *Thermodesulfobacteria* into four phyla reflecting major functional capabilities. Int J Syst Evol Microbiol 70:5972–6016.

70. Fincker M, Huber JA, Orphan VJ, Rappe MS, Teske A, Spormann AM. 2020. Metabolic strategies of marine subseafloor *Chloroflexi* inferred from genome reconstructions. Environ Microbiol 22:3188–3204.

71. l’Haridon S, Miroshnichenko M, Kostrikina N, Tindall B, Spring S, Schumann P, Stackebrandt E, Bonch-Osmolovskaya E, Jeanthon C. 2006. *Vulcanibacillus modesticaldus* gen. nov., sp. nov., a strictly anaerobic, nitrate-reducing bacterium from deep-sea hydrothermal vents. Int J Syst Evol Microbiol 56:1047–1053.

72. Kube M, Beck A, Zinder SH, Kuhl H, Reinhardt R, Adrian L. 2005. Genome sequence of the chlorinated compound–respiring bacterium *Dehalococcoides* species strain CBDB1. Nat Biotechnol 23:1269–1273.

73. Seshadri R, Adrian L, Fouts DE, Eisen JA, Phillippy AM, Methe BA, Ward NL, Nelson WC, Deboy RT, Khouri HM. 2005. Genome sequence of the PCE-dechlorinating bacterium *Dehalococcoides ethenogenes*. Science 307:105–108.

74. Peng P, Goris T, Lu Y, Nijsse B, Burrichter A, Schleheck D, Koehorst JJ, Liu J, Sipkema D, Sinninghe DJS, Stams AJM, Häggblom MM, Smidt H, Atashgahi S. 2020. Organohalide-respiring *Desulfoluna* species isolated from marine environments. ISME J 14:815–827.

75. Lie TJ, Clawson ML, Godchaux W, Leadbetter ER. 1999. Sulfidogenesis from 2-aminoethanesulfonate (taurine) fermentation by a morphologically unusual sulfate-reducing bacterium, Desulforhopalus singaporensis sp. nov. Appl Environ Microbiol 65:3328–3334.

76. Magnuson J, Stern R, Gossett J, Zinder S, Burris D. 1998. Reductive dechlorination of tetrachloroethene to ethene by a two-component enzyme pathway. Appl Environ Microbiol 64:1270–1275.

77. Fung JM, Morris RM, Adrian L, Zinder SH. 2007. Expression of reductive dehalogenase genes in *Dehalococcoides ethenogenes* strain 195 growing on tetrachloroethene, trichloroethene, or 2, 3-dichlorophenol. Appl Environ Microbiol 73:4439–4445.

78. Atashgahi S. 2019. Discovered by genomics: putative reductive dehalogenases with N-terminus transmembrane helixes. FEMS Microbiol Ecol 95:fiz048.

79. Low A, Shen Z, Cheng D, Rogers MJ, Lee PK, He J. 2015. A comparative genomics and reductive dehalogenase gene transcription study of two chloroethene-respiring bacteria, *Dehalococcoides mccartyi* strains MB and 11a. Sci Rep 5:1–12.

80. Goris T, Schenz B, Zimmermann J, Lemos M, Hackermüller J, Schubert T, Diekert G. 2017. The complete genome of the tetrachloroethene-respiring *Epsilonproteobacterium Sulfurospirillum halorespirans*. J Biotechnol 255:33–36.

81. Goris T, Schubert T, Gadkari J, Wubet T, Tarkka M, Buscot F, Adrian L, Diekert G. 2014. Insights into organohalide respiration and the versatile catabolism of *Sulfurospirillum multivorans* gained from comparative genomics and physiological studies. Environ Microbiol 16:3562–80.

82. Smidt H, van Leest M, van der Oost J, de Vos WM. 2000. Transcriptional regulation of the cpr gene cluster in ortho-chlorophenol-respiring *Desulfitobacterium dehalogenans*. J Bacteriol 182:5683–5691.

83. Kublik A, Deobald D, Hartwig S, Schiffmann CL, Andrades A, von Bergen M, Sawers RG, Adrian L. 2016. Identification of a multi-protein reductive dehalogenase complex in *Dehalococcoides mccartyi* strain CBDB1 suggests a protein-dependent respiratory electron transport chain obviating quinone involvement. Environ Microbiol 18:3044–56.

84. Mazon H, Gabor K, Leys D, Heck AJ, Van der Oost J, Van den Heuvel RH. 2007. Transcriptional activation by CprK1 is regulated by protein structural changes induced by effector binding and redox state. J Biol Chem 282:11281–90.

85. Trapnell C, Roberts A, Goff L, Pertea G, Kim D, Kelley DR, Pimentel H, Salzberg SL, Rinn JL, Pachter L. 2012. Differential gene and transcript expression analysis of RNA-seq experiments with TopHat and Cufflinks. Nat Protoc 7:562–578.

